# Transfer from number to size reveals abstract coding of magnitude in honeybees

**DOI:** 10.1101/2019.12.23.887281

**Authors:** Maria Bortot, Gionata Stancher, Giorgio Vallortigara

**Affiliations:** Center for Mind/Brain Sciences, University of Trento, 38068 Rovereto, Italy; Rovereto Civic Museum Foundation, 38068 Rovereto, Italy

**Keywords:** quantity representation, numerical cognition, magnitude, space, time and number association, honeybees

## Abstract

Number discrimination has been documented in honeybees. It is not known, however, whether it reflects, as in vertebrates, the operating of an underlying general magnitude system that estimates quantities irrespective of dimensions (e.g., number, space, time) and format (discrete, continuous). We investigated whether bees spontaneously transfer discrete discrimination of number to continuous discrimination of size. Bees were trained to discriminate between different numerical comparisons having either a 0.5 (2 *vs.* 4 and 4 *vs.* 8) or 0.67 ratio (2 *vs.* 3 and 4 *vs.* 6). Half of the subjects learnt to choose the smaller quantity and the other half the larger quantity. Bees were then tested for spontaneous choice (in the absence of reward) using comparisons with identical numbers but different sizes. Irrespective of the ratio of the stimuli, bees trained to select the smaller numerical quantity chose the congruent smaller size; bees trained to choose the larger numerical quantity chose the congruent larger size. This finding provides the first evidence for a cross-dimensional transfer between discrete (numerical) and continuous (spatial) dimensions in an invertebrate species and supports the hypothesis of a cognitive universality of a coding for general magnitude.

## Introduction

Honeybees (*Apis mellifera*) have been shown to be able to process the numerical attributes of visual stimuli (1-4), including the zero as a quantity (5). Given their distant phylogenetic origins, it is unclear, however, to what extent bees (invertebrates) and vertebrates share similarities in number cognition.

Humans and others non-human vertebrates make use of a nonverbal, nonsymbolic representation of number, the so-called Approximate Number System (ANS). The ANS obeys Weber’s law – it is thus mainly limited by the ratio between the numerical values being compared - and is thought to be supported by an evolutionarily ancient mechanism for representing quantity in an analog fashion. Gallistel (1989) first argued that discrete countable quantity (i.e., number) and continuous quantity (e.g., space and time) must be represented by a common mental currency to enable animals to perform arithmetic operations across domains (as in the case of the rate of return to a food patch, that can be computed only if organisms represent time and number in a single currency). According to this hypothesis, quantity representations in the various domains, (i.e., number, space and time), would be processed by a «common magnitude system», which represents these dimensions via the same unit of magnitude (6). Evidence that the temporal, spatial, and numerical features of a stimulus can interact with one another has been provided for vertebrates such as monkeys (7) and birds (8), and for prelinguistic human babies (9).

Interestingly, honeybees have been shown to exhibit the numerical distance effect (i.e., the fact that the ability to discriminate between numbers improves as the numerical distance increases (e.g., zero *vs.* four is easier than zero *vs.* one, (5)). The numerical distance effect is one of the signatures of the ANS and suggests the existence of an analog magnitude system in honeybees that would allow the processing of different numbers. However, it is not known whether even in bees a common set of coding mechanisms underlies quantity manipulations in different domains.

Here we investigated whether honeybees could make a transfer from discrete (number) to continuous (size) magnitudes. Bees were trained to discriminate between different numerical comparisons having either a 0.5 ratio (2 *vs.* 4 and 4 *vs.* 8) or 0.67 ratio (2 *vs.* 3 and 4 *vs.* 6). Half of the subjects learnt to choose the smaller quantity, and the other half the larger quantity. Then at test, bees were presented with stimuli of different size but identical numerosity under extinction condition (i.e., in the absence of reward). If bees possess a common mechanism to process different magnitudes, then animals trained to choose the smaller/larger quantity in the number comparisons were expected to choose the congruent smaller/larger size in the size comparison. Moreover, choice of the congruent size would not be affected by the ratio of the stimuli (i.e., ratios that proved to be discriminable for numbers should prove discriminable for sizes as well).

## Results

Honeybees proved to be able to choose the smaller number in the *number learning* test (*t*_*(15)*_ = 29.795, *P* < 0.001; Fig. 1a), and showed a significant preference for the smaller size in the *size transfer* test (*t*_*(15)*_ = 20.407, *P* < 0.001; Fig. 1b). Similarly, honeybees proved to be able to choose the larger number in the *number learning* test (*t*_*(15)*_ = 13.406, *P* < 0.001; Fig. 1a), and showed a significant preference for the larger size in the *size transfer* test (*t*_*(15)*_ = 18.331, *P* < 0.001; Fig. 1b). As in previous studies (5) we did not find any significant effect associated with the ratio, the numerical comparisons and the type of training (smaller or larger numerosity as positive) (*Number learning* test: ratio: *F*_*(1, 24)*_ = 0.432, *P* = 0.517; type of training: *F*_*(1,24)*_ = 3.860, *P* = 0.061; numerical comparison: *F*_(3,24)_ = 0.414, *P =* 0.745; ratio x type of training: *F*_*(1, 24)*_ = 0.639, *P* = 0.431; ratio x numerical comparison: *F*_*(2, 24)*_ = 0.405, *P* = 0.671; numerical comparisons x type of training: *F*_*(3,24)*_ = 1.309, *P* = 0.294; ratio x type of training x numerical comparisons: *F*_*(2, 24)*_ = 1.644, *P* = 0.214); *Size transfer* test: ratio: *F*_*(1, 24)*_ = 1.540, *P* = 0.227; type of training: *F*_*(1, 24)*_= 0.310, *P* = 0.583; numerical comparison: *F*_*(3,24)*_ = 1.119, *P* = 0.361; ratio x type of training: *F*_*(1, 24)*_= 0.016, *P* = 0.900; ratio x numerical comparisons: *F*_*(2, 24)*_ = 0.909, *P* = 0.416; numerical comparison x type of training: *F*_*(3,24)*_ = 1.547, *P* = 0.228; ratio x type of training x numerical comparisons: *F*_*(2, 24)*_ = 2.312, *P* = 0.121).

**Fig. 1.**
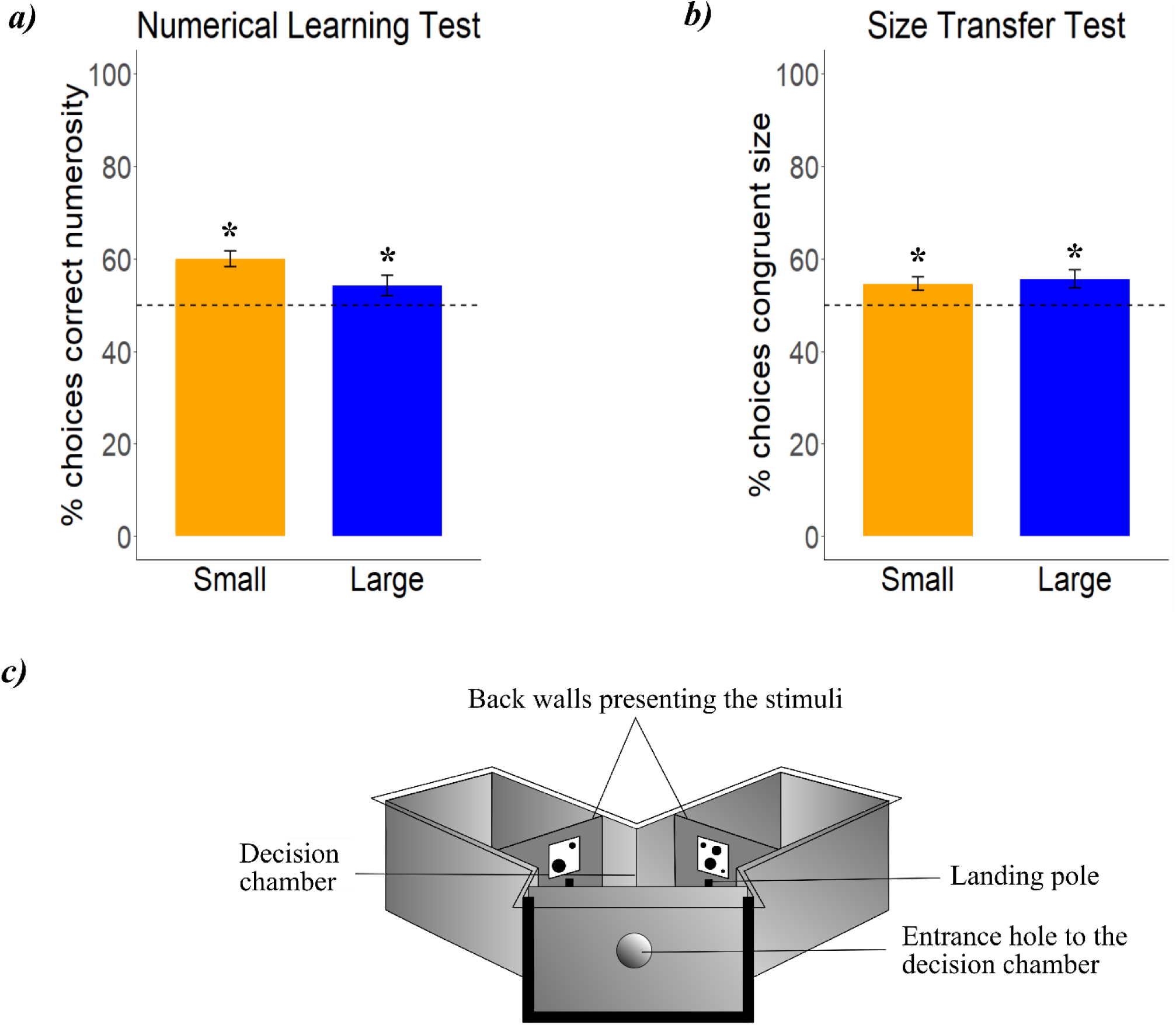
*a)* In the *number learning* test, honeybees trained to discriminate the smaller or the larger numerousness showed correct spontaneous choices in the absence of reward. *b)* In the *size transfer* test, bees showed a preference for the congruent size magnitude. (* *P* < 0.05 one-sample two-tailed *t*-tests). *c)* Schematic representation of the Y-maze used to train bees to discriminate numerousness and to test them for transfer from numerical to spatial (size) dimensions.

## Discussion

Results of *number learning* test confirmed previous studies (3-5) showing that bees can discriminate numerosities with 0.5 and 0.67 ratios when continuous physical variables were controlled for. Moreover, we found that honeybees can make a transfer from discrete (number) to continuous (size) magnitudes. This provides the first evidence for a common code for magnitudes in an invertebrate species.

The hypothesis of the existence of a prelinguistic framework to process different prothetic dimensions (i.e., dimensions that can be “more” or “less” than) was first proposed by Gallistel (6) and then developed by Walsh (10). Research in humans and other vertebrates has revealed that the temporal, spatial, and numerical features of a stimulus can interact with one another (7, 11-13) and evidence of similar activation in the parietal cortex in humans and non-human primates in quantity discrimination seems to support the hypothesis of an encoding by a common magnitude (14).

Our results show that bees generalize from a numerical dimension to a spatial (size) dimension, suggesting that a general magnitude encoding can be shared among vertebrates and invertebrates.

## Materials and Methods

Experiments were performed during the Summer 2019 at SperimentArea, a field station run by the local Natural History Museum, in Rovereto (North of Italy). Thirty-two free-flying honeybees (*Apis mellifera*) were trained singly to fly into a wooden Y-maze (fig. 1c). One half of the bees were trained with a 0.5 ratio and the other half with a 0.67 ratio. In the 0.5 ratio one half of the subjects was tested with a 2 *vs.* 4 comparison and the other half with a 4 *vs.* 8 comparison; in the 0.67 ratio one half of the subjects was tested with a 2 *vs.* 3 comparison and the other half with a 4 *vs.* 6 comparison.

The stimuli consisted of black elements, either squares, diamonds or dots on a white squared-shape background (8 cm x 8 cm) located at 15 cm distance from the decision chamber (fig. 1c). The stimuli size ranged from 1.12 cm to 3.56 cm (diameter of dots) and from 1 cm to 2.5 cm (side of squares and diamonds). The spatial disposition and the size of the elements were varied among trials to prevent the use of non-numerical cues. The continuous variables that may covary with numerosity (e.g., area, contour length, density) were varied among trials. In particular, within each shape, in one quarter of the stimuli the cumulative surface area was matched to 100%, whereas in the second quarter was not controlled (i.e., the ratio between the cumulative surface area within each pair was congruent with the numerical ratio: 0.5 in 2 *vs.* 4 and 4 *vs.* 8; 0.67 in 2 *vs.* 3 and 4 *vs.* 6). Furthermore, in the third and fourth quarter of the stimuli, the cumulative contour length was matched to 100% and not controlled, respectively, following the same logic. Additionally, half of the overall stimuli was controlled for the convex hull and the other half was controlled for the density of the elements. During the training phase, half of the bees (N=16) were presented with squares and diamonds, whereas the other half (N=16) was presented with diamonds and dots. The stimuli used in the *number learning* test, were taken from the training sample of stimuli with the area matched to 100%. In the *size transfer* test, stimuli consisted of two pairs of novel shapes (i.e., the shape that was not presented in the training phase) having sizes that differed by a ratio of either 0.5 or 0.67, depending on the numerical training previously completed by each subject. Within each pair, the two arrays had the same number and disposition of elements. In particular, the number of elements presented was equal to the numerosity reinforced during the training phase (e.g., bees trained to select 2 elements over 4 elements during the training phase, were then presented with a 2 *vs*. 2 comparison where one group of 2 elements had the double size of the other group of 2 elements).

The experimental procedure comprised a pre-training phase followed by a training and tests phase. All the phases were completed by all subjects in 1 or 2 consecutive days. During the pre-training phase, each bee was individually habituated to fly inside the apparatus and to collect food by landing on two grey poles placed in both arms, in the absence of visual stimuli. In the training phase, four different numerical comparisons (ratio 0.5: 2 *vs.* 4, 4 *vs.* 8; ratio 0.67: 2 *vs.* 3, 4 *vs.* 6) were presented to each independent group, separately. Within each group, half of the subjects was trained to select the smaller numerosity in the comparisons (either 2 or 4), whereas the other half was trained to choose the larger numerosity in the comparison (either 3, 4, 6 or 8) in order to get the food reward. During this phase, an appetitive-aversive conditioning paradigm was used: the correct numerosity was always associated with the food (0.88 M of sucrose solution) whereas the incorrect numerosity was always associated with a bitter 60 mM quinine solution, used as punishment. Each subject had to complete 60 consecutive trials of training. The stimuli were presented in a pseudo-random sequence (i.e., the correct/incorrect stimulus was never presented for more than two consecutive times on the same side).

Once completed the training phase, honeybees started the test phase. During this phase, two non-reinforced tests were presented: a *number learning* test and a *size transfer* test. Each test was presented twice to counterbalance the position of the correct array and avoid side preferences. The tests lasted 1 minute during which the number of choices (i.e., direct contact made with a body part, either the antennae or legs, on one of the two grey poles placed in front of each stimulus) made by the subjects were counted. In the *number learning* test, bees were presented with the same numerical comparisons and shapes used during the training but in the absence of any reward. In the *size transfer* test bees were exposed to the novel stimuli displaying only the size information (even in this case without any reward).

In the test phase, the percentage of correct choices was calculated for each subject and analyzed, giving rise one single value per bee to exclude pseudo-replication. All data were transformed with the arcsin transformation for proportions and percentages. The data of the *number learning* test and *size transfer* test were checked for normality (*number learning* test: Shapiro-Wilk normality test: *W* = 0.982, *P* = 0.867; *size transfer* test: Shapiro-Wilk normality test: *W* = 0.982, *P* = 0.855) and homoscedasticity (Levene’s Test on the proportion of correct choices in the *number learning* and *size transfer* test: *F*_*(1,62)*_ = 0.75, *P* = 0.389) and then analyzed with parametric statistical tests. An analysis of variance was performed with ratio (0.5 and 0.67) and type of training (smaller *vs.* larger as positive) as factors. The effect of the numerical comparisons was also analyzed with a nested factorial Anova. One-sample *t*-tests were used to assess departures from chance level (50%) in the proportions of correct choices.

## Acknowledgments

We thank Alvis Kalarikkan for his help with data collection. This project has received funding from the European Research Council (ERC) under the European Union’s Horizon 2020 research and innovation program (grant agreement No. 833504, ERC Advanced Grant SPANUMBRA to G.V.).

## Author contributions

M.B. and G.V. designed research; M.B performed the experiments; M.B and G.V. analyzed the data; G.S. contributed materials, animals and space; M.B. and G.V. wrote the paper.

